# Environmental Volatility Shifts Visual Search from Capture to Caution

**DOI:** 10.64898/2026.05.08.723763

**Authors:** Nan Qiu, Fredrik Allenmark, Siyi Chen, Hermann J. Müller, Zhuanghua Shi

**Affiliations:** The Clinical Hospital of Chengdu Brain Science Institute, MOE Key Lab for Neuroinformation, School of Life Science and Technology, University of Electronic Science and Technology of China, Chengdu, China; Neuro-cognitive Psychology, Department of Psychology, LMU Munich, 80802, Germany

**Author notes:** **Correspondent authors:** Nan Qiu, Zhuanghua Shi.

**Keywords:** visual search, attentional capture, distractor suppression, volatility, drift-diffusion model, predictive coding

## Abstract

Real-world distractors occur in environments whose states change at different rates. We asked whether such volatility alters early attentional gating or instead changes the criterion for committing to a response. Observers performed an additional-singleton search task with concurrent eye tracking while distractor presence followed high- or low-volatility sequences, with overall distractor prevalence held constant. Trial-pooled oculomotor capture was higher under high volatility, a pattern that appears to indicate altered filtering. That inference did not survive repetition-aware analysis: once the same-location run position was matched, capture did not detectably differ across volatility regimes. The pooled capture effect was therefore consistent with a structural consequence of the volatility manipulation, which enriched high-volatility blocks with early-run positions where capture is intrinsically high. The positive volatility signature appeared on distractor-absent trials, where high-volatility blocks were associated with longer target latency, more fixations, longer final-target dwell, and fewer errors. Same-location repetition learning showed no detectable difference in slope across regimes. A hierarchical drift-diffusion model (DDM) and a complementary volatility Kalman-filter (VKF) dynamic-state comparison indicated that manual responses were better described by architectures that allow both boundary-related and drift-related components than by a boundary-only account. Volatility, therefore, did not show detectable evidence of impairing the local gating rule; instead, the converging evidence points to a post-selective verification/caution profile, consistent with a precision-weighted read-out of environmental uncertainty.

## Introduction

A patch of sunshine in March means very little about whether it will be sunny an hour from now, but a sunny afternoon in July almost certainly extends into the evening. Even when the overall proportion of sunny days is identical across two weather stretches, the underlying state (sun or cloud) switches at very different rates. This contrast is what cognitive scientists call volatility: the rate at which the environment’s hidden state changes over time, independent of how often any particular state occurs (Behrens et al., 2007; Piray & Daw, 2020). Humans pick up volatility and adjust how much weight to give recent experience: when the world is unstable, recent observations carry more information; when it is stable, recent fluctuations are discounted (Behrens et al., 2007; Pulcu & Browning, 2025). This adaptive principle has been formalized as a precision-weighting operation that adjusts the gain on recent versus prior estimates (Friston, 2010; Glasauer & Shi, 2022; Piray & Daw, 2020; Pulcu & Browning, 2025). Whether the same logic reaches one of the most basic visual search operations – suppressing salient distractors during search – is only beginning to be mapped. Statistical learning of distractor regularities modulates attention at multiple levels: at the priority-map level, observers learn to down-weight predictable distractor locations (Gaspelin et al., 2025; Goschy et al., 2014; Theeuwes et al., 2022; Wang & Theeuwes, 2018); when the distractor is defined in a different dimension from the target, learning can also filter the irrelevant dimension below the priority map (Sauter et al., 2018; Zhang et al., 2019). In a previous study employing a paradigm with a long-term distractor hotspot, we showed that the degree to which observers applied that learned spatial prior for target detection depended on environmental volatility (Qiu et al., 2024). What remains to be tested is whether volatility also modulates suppression when no long-term prior exists, and suppression must be built online from short-term local repetition alone.

Distractor suppression can operate at different stages of processing. A volatile environment could change the gain of early distractor signals, the time needed to disengage after capture, and/or the criterion for committing to a response after the target has been selected. At the *pre-selective* stage, observers can down-modulate the weight of distractor signals on the search-guiding priority map before search even begins, indexed by the proportion of first saccades that land on the distractor as well as pre-display neural markers (Goschy et al., 2014; Sauter et al., 2018; Wang & Theeuwes, 2018). Convergent neural evidence indicates that the suppressive bias is genuinely prepared in advance: V1 BOLD attenuates at high-probability distractor locations (Zhang et al., 2022), and pre-array alpha lateralization predicts the amplitude of the later distractor-positivity (Pd) response (Duncan et al., 2025). At the *post-selective* stage, once observers have selected the target, what varies is the evidence required for response commitment, expressed in target dwell, fixation count, and error rate. This final stage is criterion-driven and dissociable from selection itself (Allenmark et al., 2024; Allenmark, Shi, et al., 2021), as shown by individual-differences work in which the priors used to decide *where to look* and those to decide *what is there* doubly dissociate; for instance, autistic adults show typical spatial-attention priors but atypical decision-criterion priors (Allenmark, Shi, et al., 2021). Recent proposals for terminology support the treatment of these stages as functionally distinct (Liesefeld et al., 2024). This stage logic turns the volatility question into a concrete mechanistic test: an early-gating account predicts a volatility effect on oculomotor capture after same-location repetition is matched, whereas a response-caution account predicts its clearest signature on distractor-absent trials, where target selection and response commitment can be measured without concurrent distractor-location repetition.

The standard reaction-time-based interference index (RT on distractor-present minus RT on distractor-absent trials) collapses across these stages. Worse, stage effects can move in *opposite* directions and produce a misleading null at the level of the manual response. This problem is general: the field has long recognized that pre-selective gating and rapid reactive components are difficult to dissociate from RT alone, motivating EEG- and eye-tracking-based decompositions (Liesefeld et al., 2017; McDonald et al., 2025). Bogaerts et al. (2022) and van Moorselaar and Slagter (2019) showed that manipulating distractor regularities over different timescales modulates suppression, suggesting that suppression is sensitive to the structure of distractor regularities. However, these studies do not isolate *volatility*, the rate at which the underlying state switches with the base rate held constant, as a distinct manipulation. Our recent work began to address this gap by crossing volatility with a long-term spatial bias, showing that volatility interacts with the deployment of a learned spatial prior in target detection (Qiu et al., 2024). Extending that test requires both a design that eliminates the spatial probability prior and an analytical pipeline that separates pre-selective gating, engagement/disengagement, and post-selective verification, so that any dissociation between them can be expressly measured rather than assumed.

We addressed that gap using an equal-probability eight-location search display. When present, the distractor appeared at each non-target location equally often and repeated at the same location across consecutive distractor-present trials until a streak broke. Volatility was manipulated across two within-subjects sessions by varying the Markov transition probability governing state persistence (*p* = 0.3 vs. *p* = 0.7), while overall distractor prevalence was maintained at 50%. This eliminates any long-term spatial hotspot, but the transition rule that defines volatility necessarily produces different mixtures of repetition streaks across regimes, so a pooled present-trial contrast cannot by itself separate a per-trial volatility effect from a repetition-mixture effect. We treat this property as a design feature and complement the pooled contrast with two repetition-aware analyses: matched-repetition slices that hold the same-location repetition constant across regimes, and distractor-absent trials that probe target verification and response commitment without concurrent distractor competition.

In this study, we asked three questions. First, does high volatility produce higher oculomotor capture even when the same-location repetition is matched, as an early-gating account predicts? Second, does high volatility instead appear most clearly in distractor-absent measures of target verification and response commitment, as a post-selective caution account predicts? Third, does the local learning rule that converts same-location repetition into suppression change its slope across volatility regimes, or is the repetition-learning rate preserved while the decision state shifts? We complement these repetition-aware behavioral and eye-tracking tests with two manual-response models. A hierarchical DDM (Fengler et al., 2022; Ratcliff & McKoon, 2008) provides a condition-level latent-decision decomposition of drift, boundary, and non-decision time, and a reduced Volatile Kalman Filter (VKF) comparison asks whether trial-wise latent volatility/prediction-error state improves RT prediction through drift-like modulation, boundary-like modulation, or both. The mechanistic inference rests on the joint pattern: repetition-aware ocular measures constrain the gating claim, the distractor-absent family supplies the positive stage-specific signature, and the modelling clarifies which decision-state axes best describe manual responses.

## Methods

### Participants

We recruited 24 healthy adults (age range: 18-40 years; mean age = 26.5 years; 12 females and 12 males) for two experimental sessions (high- and low-volatility). For the volatility manipulation, the sample size was estimated based on the high-vs. low-volatility effect reported by Qiu et al. (2024), corresponding to a within-participant effect size of *dz* = 0.73. A two-sided paired-samples calculation with alpha = 0.05 and power = 0.80 yielded a required sample size of 17 participants. We increased the sample to 24 to account for uncertainties in eye movements. After exclusions for incomplete eye-tracking sessions (see

*Awareness assessment*, below), the final analysis sample was *N* = 21; a sensitivity analysis indicates that this sample retains 80% power at α = 0.05 (two-sided) to detect within-subject paired effects of *d*z ≥ 0.64. All participants reported normal or corrected-to-normal vision. They provided informed consent and received course credits or financial reimbursement (9 € per hour). The study was approved by the ethics committee of the Department of Psychology and Pedagogics at LMU Munich.

### Apparatus

Testing took place inside a dimly lit, sound-attenuated, electrically shielded cabin. An EyeLink 1000 desktop-mounted eye tracker (SR Research, Canada) sampled the dominant eye at 1 kHz, while a 24-inch VIEWPixx/3D monitor (VPixx Technologies Inc.; 1920×1080 pixels, 120 Hz) rendered the search displays. A custom MATLAB R2019b script, built on Psychtoolbox and the EyeLink Toolbox (Brainard, 1997), controlled stimulus generation, response logging, and gaze sampling and routed keyboard input to the trial loop. Throughout each session, a chin rest held participants at approximately 65 cm from the screen.

### Stimuli

Each search display arrayed 40 grey bars (0.18 × 0.81°) on a black background (RGB = 0 0 0), spaced around four concentric rings centered on the fixation cross, with radii of 1.1°, 2.2°, 3.3°, and 4.4°. Every bar carried a small notch (∼0.25° tall) in either its upper or lower half, with notch position randomized and balanced within each block (Fig 1A). The bulk of these bars formed a homogeneous background of vertical (0°) elements rendered at 20% of maximum luminance. Embedded within that background, the target tilted 12° to the right at the same intensity, while the orientation distractor, when present, tilted 45° to the left, carried two notches, and matched the non-target intensity. Critically, target and distractor locations alike were drawn from the same set of eight positions on the second concentric ring (Fig 1A).

**Fig 1.**
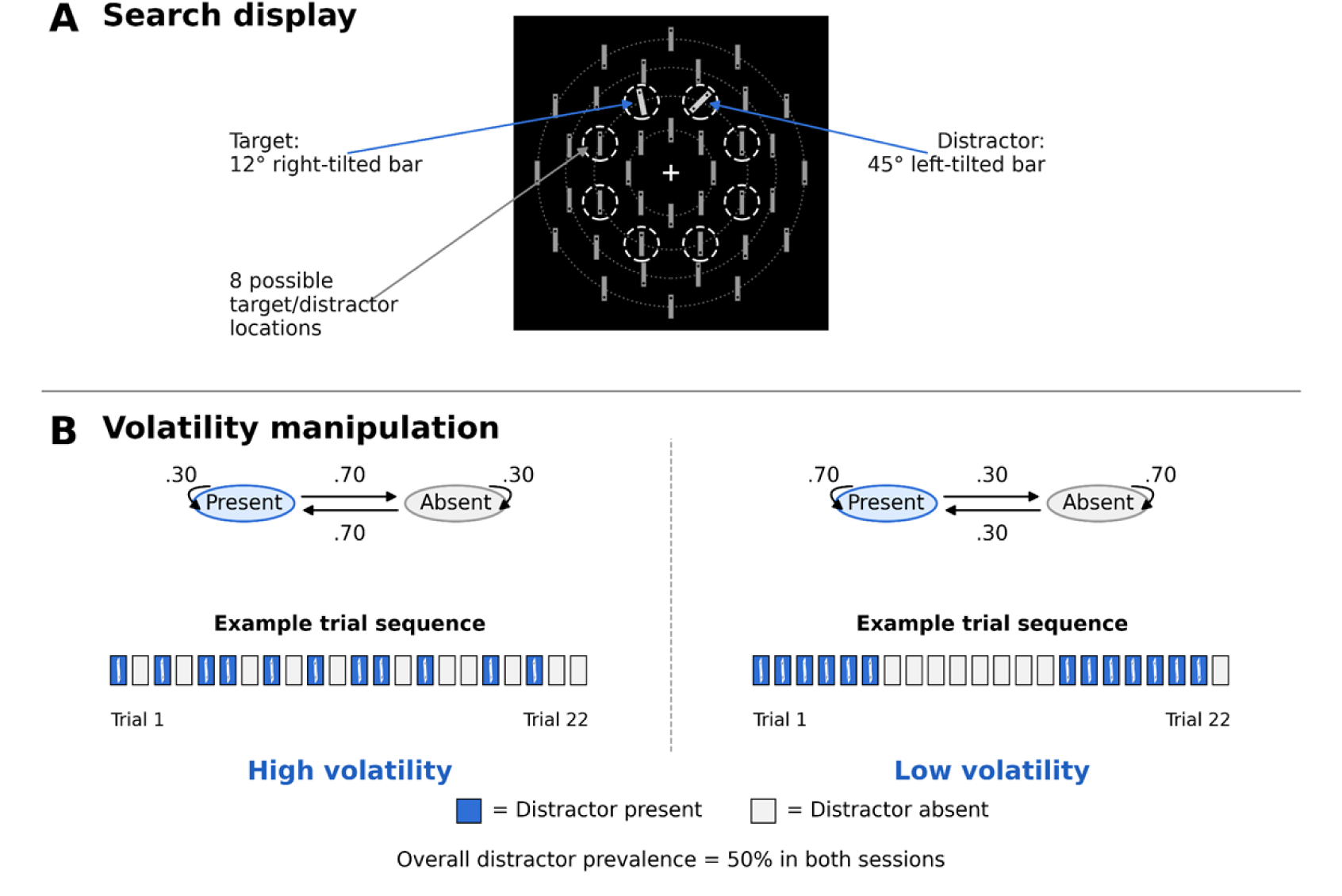
Search display and volatility manipulation. (**A)** Observers searched for a 12°-tilted target bar and reported the position of its notch (top or bottom). In some trials, a task-irrelevant 45°-tilted orientation distractor also appeared and was to be ignored. Target and distractor locations were drawn from the same set of eight equally spaced positions on the second concentric ring. (**B)** A two-state Markov process over distractor presence (Present ↔ Absent) defined the volatility regime, with transition probabilities relating trial *n* to trial *n*−1 (PP, PA, AP, AA). In the high-volatility session (left), the state switched more often than it repeated (PP = AA = 0.30; PA = AP = 0.70); in the low-volatility session (right), it mostly repeated (PP = AA = 0.70; PA = AP = 0.30). Example 22-trial sequences below each diagram illustrate the resulting structure (filled square = distractor present; empty square = distractor absent). Overall distractor prevalence remained at 50% in both sessions, with participants completing 5–7 days apart. *Note:* PP = distractor present following present; PA = distractor absent following present; AP = distractor present following absent; AA = distractor absent following absent.

### Design

We generated each participant’s trial sequence using the *Markov-chain* package in R, with a two-state Markov process over distractor presence (Present/Absent). Both sessions drew from the same generative scaffold: 1440 analyzed trials, the two states equally frequent (1/2 each), overall distractor prevalence fixed at 50%, and a unique sequence drawn for every participant, which differed only in the transition matrix relating trial *n* to trial *n*−1. The high-volatility session made switches more likely than repeats (PP = AA = 0.30; PA = AP = 0.70), producing rapid alternation; the low-volatility session reversed this structure (PP = AA = 0.70; PA = AP = 0.30), yielding longer runs of the same state (Fig 1B). Each session opened with 80 unanalyzed practice trials and continued with 9 blocks of 160 analyzed trials.

In every distractor-present run, the distractor was repeated at the same location across consecutive trials. Because random sequences vary in run-length distribution, we screened candidate sequences to match the count of three-trial distractor-repeat subsequences across the two volatility conditions. Three-trial repeats are the longest repetition we analyzed and the level with the thinnest statistical power, so matching them between regimes is a precondition for any volatility-by-repetition contrast. Retained high-volatility sequences contained 31–39 such subsequences, and low-volatility sequences contained 31–51 sequences, both anchored at the same 31-subsequence floor (Fig 1B).

Distractor locations were not globally biased across the experiment. At the onset of each distractor-present run, the location was drawn from the eight possible target/distractor positions; subsequent trials within the run reused that location. The design therefore isolates short-term repetition-based suppression within runs rather than long-term spatial-probability learning of any high-probability location.

### Procedure

Participants performed a classic visual search task. On each trial, they searched for the target (a bar tilted 12° to the right) and indicated the notch orientation (top or bottom) by pressing a mouse button with either their right or left index finger. They were instructed to fixate on the cross as soon as possible after finding the target.

Each trial began with a fixation cross presented for a random interval of 0.8 to 1.6 s. The visual search display remained visible until a response was provided or for up to 4 s. Participants were instructed to respond as quickly as possible without sacrificing accuracy. After an incorrect or delayed response, the fixation cross changed color for 1 s: red for an incorrect response and blue for a slow response. Inter-trial intervals contained the fixation cross and were jittered randomly from 0.8 to 1.6s before the next trial.

After completing both sessions, participants answered an awareness probe: *“Do you think the irrelevant tilted-left 45-degree bar changes its location more frequently from trial to trial?”*, choosing one of the following: (1) more often in Session 2 than Session 1; (2) more often in Session 1 than Session 2; (3) the location did not change obviously in either session; (4) the location frequently changed in either session; or (5) unsure. Eleven participants chose “unsure”, three each chose options (3) and (4), and seven chose option (1) or (2); of these seven, only one answered correctly. Three participants (IDs 7, 14, and 17) were excluded from all analyses because the eye-tracking record for one of their two volatility sessions was incomplete, leaving the within-subject high-vs-low contrast, on which all reported analyses depend, incomputable. This exclusion was fixed on data-completeness grounds before any awareness frequencies were tabulated and is therefore independent of awareness; the one participant who answered correctly happened to be among the three excluded.

### Computational models

A hierarchical Bayesian drift-diffusion model using HSSM 0.3.0 (Fengler et al., 2022) was used to assess whether the manual-response profile is better described by changes in drift rate, decision boundary, or both. The HSSM has four key parameters: drift rate *v*, boundary *a*, non-decision time *t*, and initial bias *z*. Given that the response features (top vs. bottom notch on the target) is unbiased and orthogonal to the volatility manipulation, *z* was not modeled here. Four nested specifications were compared: M0, with no condition effects; M1, in which drift varied with Volatility × Distractor; M2, in which boundary varied with Volatility; and M3, in which drift varied with Volatility × Distractor while boundary and non-decision time varied with Volatility. Full priors, posterior-predictive checks, and identifiability caveats are reported in Supplementary S1.

As a complementary dynamic-state analysis, we fit reduced volatility Kalman-filter (Piray & Daw, 2021) + DDM-style RT models that linked a trial-wise latent volatility/prediction-error state to drift-like modulation, boundary-like modulation, or both. Four subject-wise architectures were compared: *Null (gain-only)*, in which distractor-present gain affected drift with no latent-state modulation; *Drift-only (M1)*, in which the latent VKF state modulated drift; *Boundary-only (M2)*, in which the latent VKF state modulated boundary; and *Full (M3)*, in which the latent VKF state modulated both drift and boundary. The observation model combined Bernoulli correctness with Gaussian RT around analytic DDM mean moments, and subject-wise fits were aggregated with AIC/BIC. Because this model is a simplified dynamic-state approximation rather than a hierarchical Wiener likelihood, it is treated as a supportive architecture check rather than as the primary modelling analysis.

## Results

### Overview of experiments

The volatility manipulation is implemented through a Markov transition probability, which means it can alter behavior through two non-exclusive routes. A *structural* route reaches selective attention through the run-position mixture each regime imposes on the trial sequence: the local same-location repetition-learning rule, fed a different mixture of trial histories, produces a different mean phenotype. A *modulatory* route is a per-trial change in gain or decision criterion on top of that mixture. We therefore organize the results such that each route can be evaluated separately and at the processing stages predicted in the introduction. The first set of analyses reports the trial-pooled signatures of the manipulation, which by construction sum the two routes. The second set isolates the per-trial modulatory route by matching same-location repetition between regimes, and asks where in the processing stream — pre-selective gating, post-selective verification, or the local learning slope — that route leaves a detectable signature. The third set fits hierarchical-Bayesian and reduced VKF latent-decision models to manual responses, providing a latent-decision counterpart to the eye-tracking conclusion.

### The pooled phenotype: structural and modulatory routes combined

#### Trial-pooled RT, error rates, and oculomotor capture

A 2 (Distractor condition: absent vs. present) × 2 (Volatility: high vs. low) repeated-measures ANOVA on correct-trial RTs showed the expected distractor-interference effect: responses were slower when the singleton distractor was present than when it was absent (1129.7 vs. 976.0 ms), *F*(1, 20) = 178.07, *p* < 0.001, η_p_² = 0.90. The main effect of volatility was marginal, with slower responses in high-than low-volatility sessions (1107.3 vs. 998.5 ms), *F*(1, 20) = 4.15, *p* = 0.055, η_p_² = 0.17; the Distractor × Volatility interaction was not reliable, *F*(1, 20) = 0.68, *p* = 0.420, η_p_² = 0.03 (Fig 2A-B). Errors were rare overall (≈2.2%) and showed a volatility main effect, *F*(1, 20) = 6.78, *p* = 0.017, η_p_² = 0.25 (1.7% high vs. 2.7% low), with neither the distractor main effect nor the interaction reliable (*p*s > 0.397).

**Fig 2.**
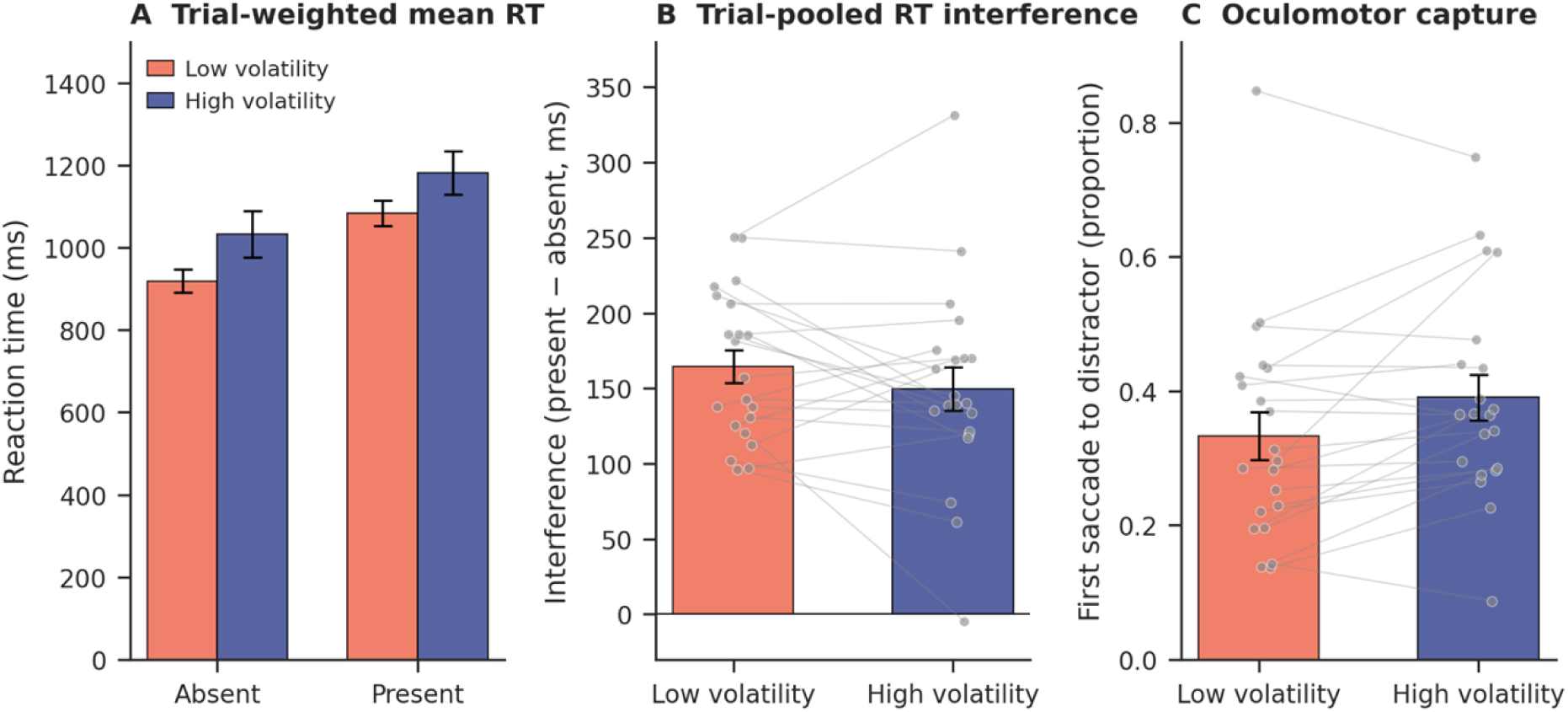
Trial-pooled signatures of the volatility manipulation. **(A)** Mean RT in the four cells of the Volatility × Distractor design; the Distractor × Volatility interaction is not reliable, *F*(1, 20) = 0.68, *p* = 0.42. **(B)** Trial-pooled per-subject RT-defined distractor interference. **(C)** Proportion of first saccades to the distractor on present trials, trial-pooled: higher with high vs. low volatility, paired *t*(20) = 4.48, *p* = 0.001. These conventional pooled summaries motivate the repetition-aware decomposition below; they are not, by themselves, per-trial mechanistic tests.

Pooling across same-location run positions, the proportion of first saccades to the distractor on present trials was higher in high-than low-volatility sessions (38.6% vs 29.8%; +9.0 percentage points, pp), paired *t*(20) = 4.48, *p* = 0.001, *dz* = 0.98 (Fig 2C). Conventionally, the pooled signature is interpreted as a per-trial volatility effect on early selection.

Taken at face value, manual RTs on distractor-present trials suggest a puzzle (Fig 2A). High-volatility contexts slowed overall responding and produced fewer errors, yet the distractor interference (RT_present_ - RT_absent_) was not reliably modulated by volatility in the omnibus 2 × 2 ANOVA above. The eye-tracking record, in contrast, already shows the *opposite* sign on its own pre-selective measure: oculomotor capture by the distractor was substantially higher in high-than in low-volatility sessions (Fig 2C).

#### Distractor-absent trials

On distractor-absent trials, five measures shifted between regimes in a coherent direction (Fig 3). RT was longer in high-than low-volatility sessions (+112.8 ms), *t*(20) = 2.08, *p* = 0.051, *d*z = +0.45; and errors were rarer (−0.91 pp), *t*(20) = −2.46, *p* = 0.023, *d*z = −0.54; saccade-to-target latency rose by +31.1 ms, *t*(20) = 2.64, *p* = 0.016, *d*z = +0.58; total fixations rose by +0.34, *t*(20) = 2.75, *p* = 0.012, *d*z = +0.60; and final target dwell rose by +96.0 ms, *t*(20) = 2.46, *p* = 0.023, *d*z = +0.54. Effect sizes were uniformly medium-small (|*d*z| = 0.45–0.60). This family shows a positive volatility signature in the dataset: high volatility was associated with slower target acquisition, prolonged verification, and fewer errors on trials without a distractor.

**Fig 3.**
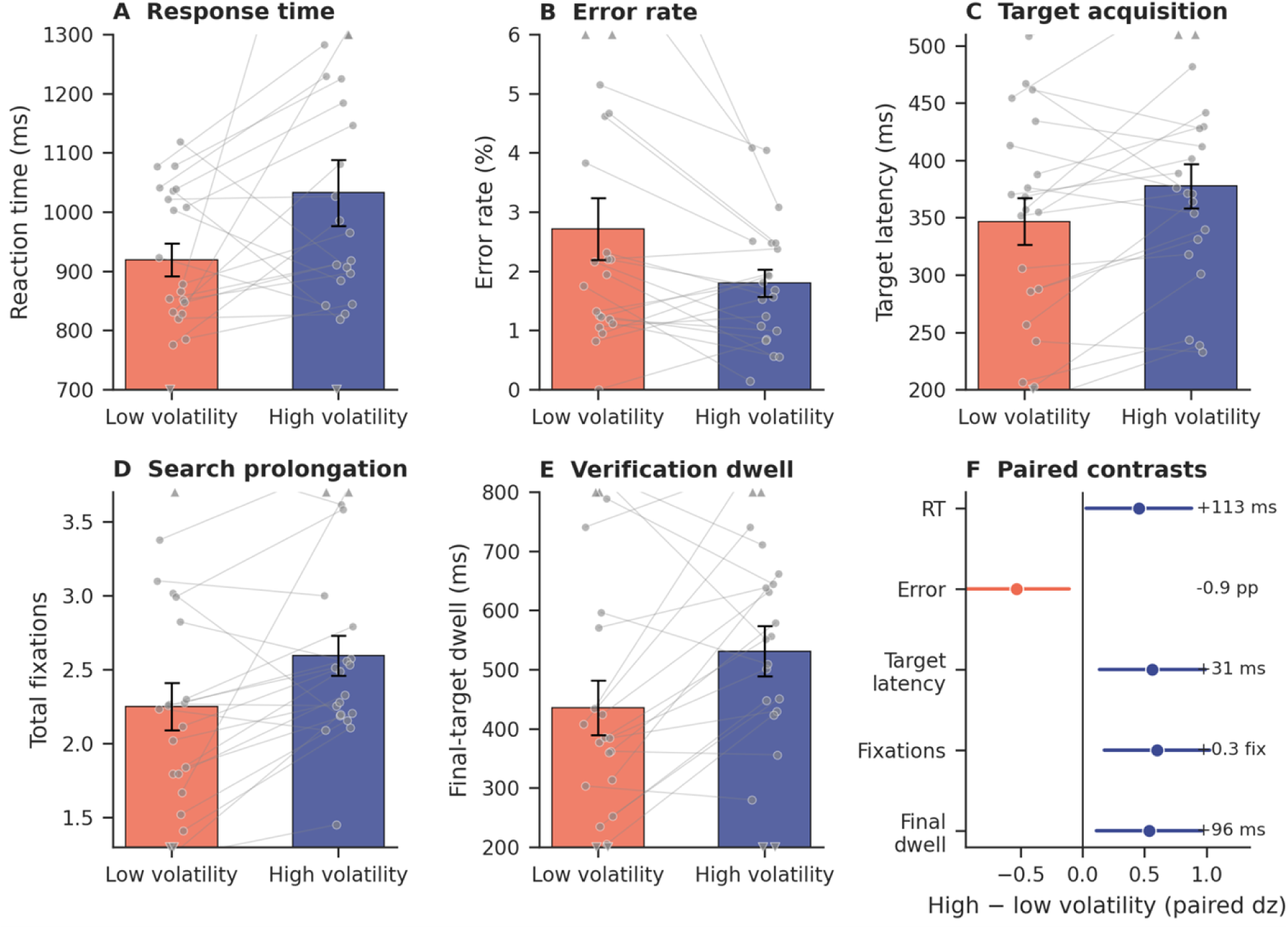
Distractor-absent volatility profile. (A-E) Paired subject summaries for the five primary distractor-absent measures: reaction time, error rate, target latency, total fixations, and final target dwell. Bars show condition means; grey lines link the same participant across volatility sessions; error bars are ±1 SEM. Subjects whose values fall outside the plotted range are drawn as small triangles at the corresponding axis edge (▴ above, ▾ below), keeping bar heights comparable across panels. **(F)** Point locations show paired high-minus-low contrasts in within-subject *d*z units; right-side labels give the corresponding raw high-minus-low differences in native units. The pattern links longer target acquisition and verification with fewer errors under high volatility.

### Isolating the modulatory route at matched repetition

The trial-pooled present-trial contrasts in the previous subsection mix three populations of trials in regime-specific proportions, while the absent-trial contrasts avoid distractor-location repetition by design. The frequency of same-location repetition run positions varies across the two volatility regimes, and several distractor-suppression measures depend strongly on run position. We therefore re-examined each present-trial signature at its matched run position and then evaluated the absent-trial family with explicit multiplicity control.

#### The repetition mixture imposed by the volatility

The two volatility regimes generate different distributions of consecutive same-location run positions on present trials. In high-volatility sessions, the distractor repetition pool is approximately 70% rep 1 / 21% rep 2 / 9% rep 3+; in low-volatility sessions, the split is 31% / 23% / 47%. Oculomotor capture is intrinsically higher at repetition 1 than at repetition 3 (combined-volatility means ≈43% vs. ≈24%), so any pooled present-trial mean compares a repetition-1-heavy mean under high volatility with a repetition-3-heavy mean under low volatility.

#### Present-trial effects at matched same-location repetition

A 2 (Volatility) × 3 (Repetition: rep 1, rep 2, rep 3+) repeated-measures ANOVA on the proportion of first saccades to the distractor returned no reliable volatility main effect, *F*(1, 20) = 0.24, *p* = 0.626, η_p_² = 0.012; there was a strong monotonic repetition decay, *F*(2, 40) = 67.49, *p* < 0.001, η_p_² = 0.77, and a non-reliable interaction, *F*(2, 40) = 1.36, *p* = 0.267 (Fig 4A). Per-rep gaps (high − low) were small and unsystematic in sign: +1.1 pp at rep 1, +2.7 pp at rep 2, −1.1 pp at rep 3+. Bayesian and frequentist equivalence tests gave a bounded null interpretation. Bayes factors favoured the null overall (rep-equal pooled BF₀₁ = 2.60), with moderate support at rep 1 (BF₀₁ = 4.11) and rep 3+ (BF₀₁ = 3.03) and only anecdotal support at rep 2 (BF₀₁ = 1.43); the corresponding TOST at SESOI *d*z = ±0.4 was inconclusive, reflecting the limited sensitivity with *n* = 21. We therefore consider capture at matched repetitions as consistent with no detectable per-trial difference.

**Fig 4.**
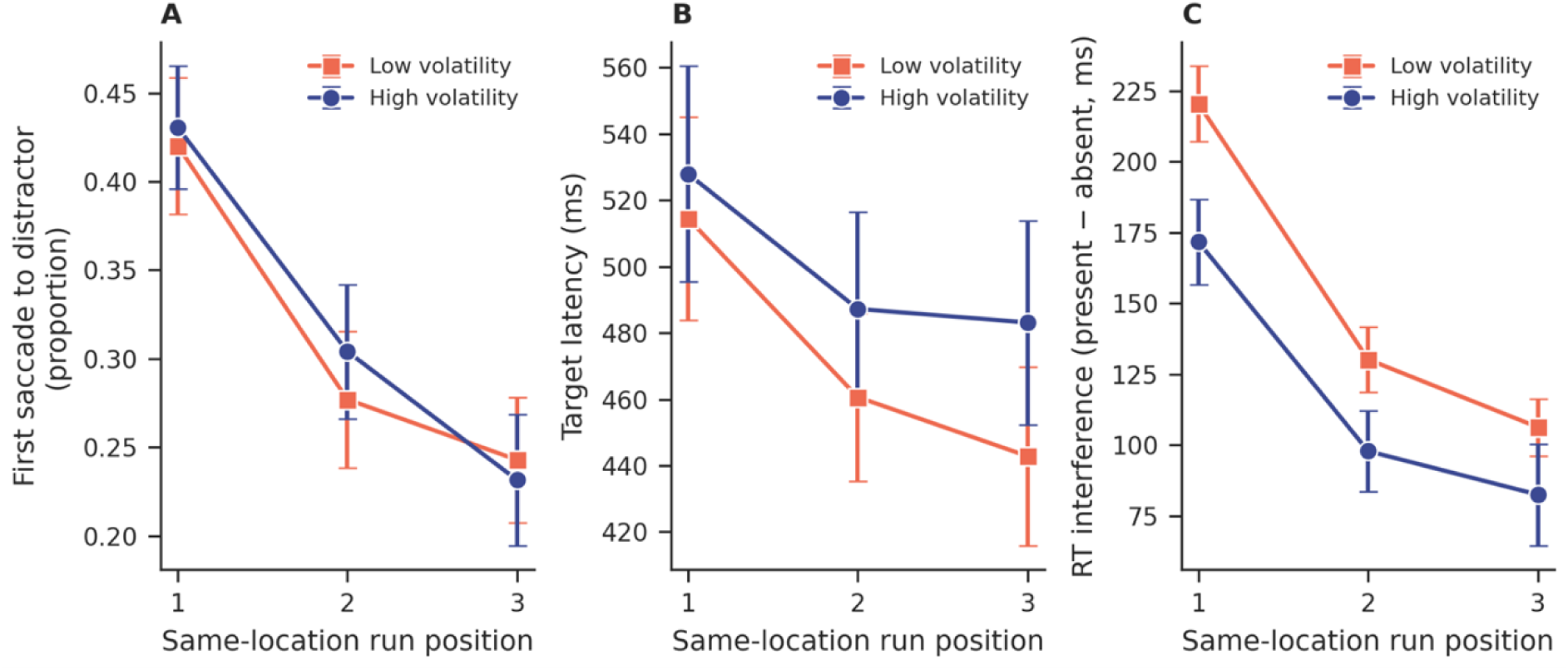
Repetition-driven learning across volatility contexts. Same-location run position on the x-axis (rep 1 = first distractor-present trial of a new run after an absent trial; rep 2–3 = consecutive same-location continuations); high-volatility (blue) and low-volatility (orange) sessions overlaid. **(A)** Proportion of first saccades directed to the distractor. **(B)** Target latency (ms). **(C)** RT interference (present − absent, ms). On every measure, repetition produced a strong monotonic decay that was statistically equivalent across volatility regimes (LMM slope-equivalence LRT *p* = 0.508, 0.320, and 0.174 for A, B, C). Points show condition means; error bars are ±1 SEM. Volatility shifts the offset of the curve without changing its per-repetition slope.

The same 2 × 3 ANOVA on present-trial RT yielded no reliable volatility main effect, *F*(1, 20) = 2.10, *p* = 0.163, η_p_² = 0.095, although per-rep gaps were directionally consistent with a slight slowing under high volatility (+65 / +81 / +90 ms across reps 1/2/3+; all *p* > 0.13). When RT-defined distractor interference was considered under repetition-equal weighting (the mean across reps 1, 2, and 3+ rather than across raw trials), the cost was smaller under high than low volatility, *F*(1, 20) = 10.22, *p* = 0.005. This repetition-equal cost is not a trial-pooled signature; it reflects the contrast between matched present-trial run positions and the absent-trial baseline. The trial-pooled present-trial signatures reported above are therefore consistent with the rep-mixture imposed by the Markov manipulation, rather than a detectable per-trial effect on capture.

#### A converging post-selective signature on distractor-absent trials

The analysis of matching same-location repetition revealed no detectable per-trial capture effect on the distractor-present trials. The five absent-trial measures reported above are repetition-unconfounded by design, but they are not five independent tests. Across-subject Δ correlations confirm tight coupling between target latency and total fixations (*r* = +0.75) and a speed–accuracy axis between error rate and final dwell (*r* = −0.46); the five-vector spans roughly two underlying axes: deployment-prolongation and a speed–accuracy tradeoff.

A multivariate Hotelling’s *T*² on the joint 5-vector of paired differences against zero was marginal, *T*² = 14.50, *F*(5, 16) = 2.32, *p* = 0.092; the latency–fixation coupling makes the joint test conservative. Marginal multiple-comparison control across the family yielded 4/5 surviving Benjamini–Hochberg FDR at *q* = 0.05 (target latency, total fixations, final-target dwell, and error rate, all at *p*_FDR_ = 0.029). The RT effect did not survive correction (*p*_FDR_ = 0.051), and we therefore interpret it as part of the broader verification profile rather than as a stand-alone effect. Read across the family, the converging direction (slower target acquisition, more fixations, longer dwell, fewer errors) is the clearest repetition-unconfounded signature of high volatility.

A 2 × 3 RM-ANOVA on each measure with distractor-absent run position showed that this signature was present from rep 1 and amplified with run position: total fixations *F*(1, 20) = 5.62, *p* = 0.028; final-target dwell *F*(1, 20) = 5.25, *p* = 0.033; target latency *F*(1, 20) = 3.84, *p* = 0.064 (high − low gaps of +14, +26, +30 ms across reps 1/2/3); first-saccade-to-target proportion *F*(1, 20) = 3.37, *p* = 0.082.

A second effect on distractor-absent trials concerns *which* object the first saccade reaches. The proportion of first saccades that landed on the target was higher under high than low volatility (52.9% vs. 47.9%; +5.0 pp), paired *t*(20) = 1.84, *p* = 0.082. Decomposed by destination, first-saccade latency on absent trials was longer under high volatility overall (+14.2 ms, *p* = 0.019), but the rise was carried by saccades to non-singleton distractors (+28.3 ms, *t*(20) = 3.07, *p* = 0.006, *d*z = +0.67) rather than by saccades to the target (−11.6 ms, *p* = 0.291). On any given trial, target latency = first-saccade latency + post-launch interval, where the post-launch interval is the time from saccade launch until the eyes first reach the target (lengthened whenever the first saccade lands on a non-target placeholder so a corrective saccade is needed). The same identity carries over to the high-minus-low contrasts, so the +31 ms volatility-related rise in target latency partitions into a +14 ms rise in first-saccade latency and a ∼17 ms rise in the post-launch interval. The across-subject coupling between Δprop-to-target and Δfirst-saccade-latency was null (*r* = +0.18, *p* = 0.425): the latency rise does not buy a corresponding rise in target-reaching success, so a “wait longer to direct better” account is not supported. This allocation component accompanies the broader verification profile, but it should not be treated as proof of a response-criterion shift on its own.

#### Same-location repetition learning is rate-invariant across regimes

Linear mixed-effects models on subject × volatility × run-position cell means tested whether volatility shifts the per-repetition learning slope rather than only the curve’s offset. On every measure, the slope-difference term was non-significant. First-saccade-to-distractor: β_int_ = −0.011, χ²(1) = 0.44, *p* = 0.508. Target latency: β_int_ = +13.4 ms/rep, χ²(1) = 0.99, *p* = 0.320. RT-defined distractor interference: β_int_ = +12.5 ms/rep, χ²(1) = 1.84, *p* = 0.174. Each measure showed a strong repetition main effect (e.g., first-saccade-to-distractor 0.43 → 0.24 across reps 1/2/3+; RT-defined cost 196 → 94 ms). We therefore found no detectable volatility-related slope difference in the local learning rule that converts same-location repetition into spatial suppression (Fig 4).

### Computational modelling of manual responses

#### Hierarchical DDM provides a descriptive drift/boundary decomposition

The manual RT/error profile was also fit with a hierarchical Bayesian drift-diffusion model (HDDM-style DDM, implemented in HSSM). Model comparison favored the full model M3 – in which drift rate varied with Volatility × Distractor and both decision boundary and non-decision time varied with Volatility – over the drift-only, boundary-only, and null models (M3 stacking weight = 0.71; Δelpd_loo relative to M3 = 383 for M1, 686 for M2, and 953 for M0; Fig 5C). This indicates that the manual-response likelihood contains information about both drift-related and boundary-related volatility effects.

**Fig 5.**
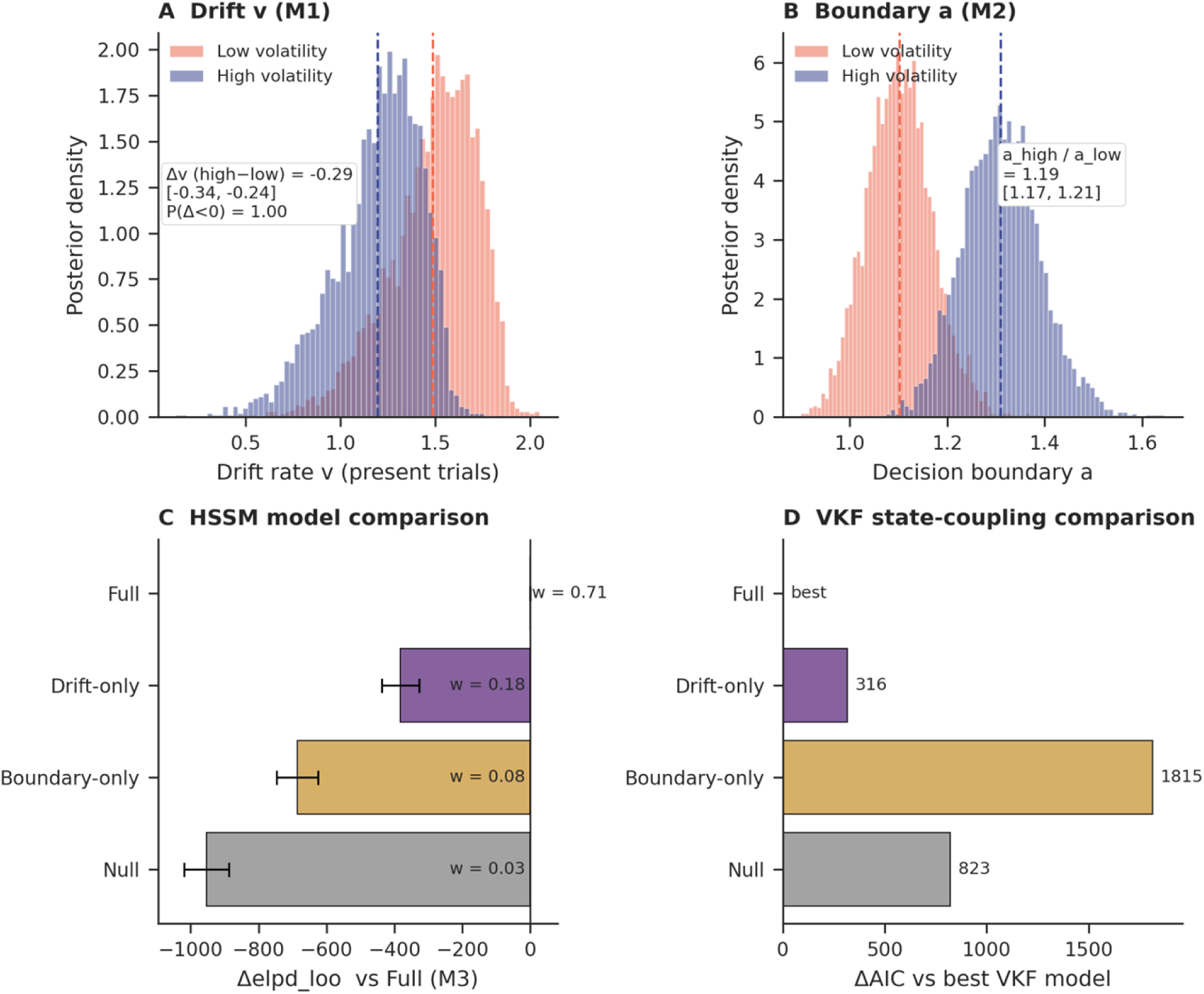
Computational modelling of manual responses. **(A)** Posterior distributions for present-trial drift rate *v* in the HSSM drift-only model M1. **(B)** Posterior distributions for decision boundary *a* in the HSSM boundary-only model M2. **(C)** HSSM PSIS-LOO model comparison, plotted as Δelpd_loo relative to the full model (M3); more negative values indicate poorer expected predictive accuracy than the full model. **(D)** Reduced VKF + DDM-style model comparison, plotted as ΔAIC relative to the best VKF model. Panels C and D use the same row order (top→bottom: Full, Drift-only, Boundary-only, Null) and the same colour scheme per architecture, so the two model families can be read against each other directly. The HSSM provides the primary latent-decision decomposition; the VKF panel provides a supportive dynamic-state architecture check, not a separate stage-specific inference.

Because the full model’s cell-level parameter magnitudes lie on a known parameter-identifiability ridge (drift rate *v, boundary a*, non-decision time *t*), we use the lower-dimensional models as interpretable projections of the two decision axes, rather than as a uniquely identified joint decomposition. In the drift-only projection (M1), present-trial drift was lower under high than low volatility, Δv (high−low) = −0.29, 95% HDI [−0.34, −0.24] (Fig 5A). In the boundary-only projection (M2), the decision boundary was higher under high volatility, *a*_high_ / *a*_low_ = 1.19, 95% HDI [1.17, 1.21] (Fig 5B). The boundary component provides a latent-decision counterpart to the distractor-absent verification/caution profile. The drift component shows that the manual RT distribution is not explained by boundary shift alone, but it should not be read as independent evidence for a detectable per-trial early-selection change: the present-trial manual drift estimates are affected by the same repetition-mixture structure that motivates the matched-repetition eye-movement analyses.

#### A reduced VKF comparison tests trialwise state coupling

A reduced VKF + DDM-style RT comparison asked a related dynamic-state question: does a trialwise latent volatility/prediction-error state improve prediction through drift-like modulation, boundary-like modulation, or both? The four VKF models are named in parallel to the HSSM specifications M0-M3 (Null, Drift-only, Boundary-only, Full). The full drift/boundary model provided the best aggregate AIC across 60,255 RT-bounded trials from 21 subjects (AIC = 80245.84), followed by the drift-state model (ΔAIC = 316.26), the gain-only baseline (ΔAIC = 822.57), and the boundary-state model (ΔAIC = 1815.18; Fig 5D). Subject-wise diagnostics showed that the joint model improved negative log likelihood relative to the gain-only baseline for 21/21 subjects (median ΔNLL = −5.93), while the drift-state and boundary-state models improved fit for 19/21 and 16/21 subjects, respectively.

The VKF comparison is deliberately treated as supportive rather than decisive. It uses a simplified Bernoulli-correctness plus Gaussian-RT likelihood and subject-wise maximum-likelihood fits, whereas the HSSM uses a hierarchical Wiener likelihood. Its AIC ranking converges with the HSSM at the architectural level: manual RT is best described when both drift-like and boundary-like modulation are allowed. BIC, however, favored the simpler gain-only baseline because the subject-wise dynamic models carry many additional parameters (BIC = 82014.08 for the gain-only vs. 82326.31 for the drift-boundary model). The VKF analysis, therefore, does not provide an independent mechanistic identification of boundary setting or early gating. It instead supports the narrower conclusion that the manual-response data are not a pure-boundary story and that dynamic state information is useful only as a secondary check on the HSSM interpretation.

## Discussion

Volatility changes how observers commit to visual decisions, while the early-selection process that gates out distractors remains intact. The data support that conclusion through coordinated answers to the three questions raised in the introduction. When consecutive same-location run position was matched between regimes, oculomotor capture showed no detectable per-trial volatility difference; the +8.8 pp pooled rise was reachable from the run-position mixture imposed by the manipulation alone, without any additional gating modulation. On distractor-absent trials, where no distractor-location repetition can confound the contrast, high volatility produced a coherent post-selective profile: slower target acquisition, prolonged verification, more fixations, and fewer errors. Manual responses were assigned to that profile with a higher latent decision boundary in the HSSM analysis. The local same-location repetition-learning slope was statistically indistinguishable across regimes, indicating that the rule that converts repeated same-location distractors into spatial suppression operates the same way in volatile as in stable environments. Together, volatility acts on visual selection through two complementary routes: a *structural* route arises because the same local suppression rule, fed a different mixture of trial histories, produces a different mean phenotype. A *modulatory* route adds a per-trial shift in decision criterion on top of that mixture. In this design, the structural route accounts for the effect of volatility on early selection, while the modulatory route carries its post-selective signature.

The modulatory route maps directly onto a precision-weighting account of volatility. In adaptive frameworks, volatile environments lower confidence in long-run state estimates and raise the weight assigned to recent observations (Behrens et al., 2007; C. Mathys et al., 2011; Piray & Daw, 2020; Pulcu & Browning, 2025; Yu & Dayan, 2005). A natural read-out of that uncertainty is a higher response criterion: when the current state is less certain, observers require more confirmatory evidence before committing (Forstmann et al., 2016). The distractor-absent profile fits that prediction directly: target acquisition and dwell were prolonged, fixations increased, and errors decreased. The boundary-only HSSM projection (*a*_high_ / *a*_low_ = 1.19) supplies a formal latent-decision counterpart in the manual response. Pre-selective gating operates on the current display, and once the same-location repetition is matched, it reveals no detectable volatility-dependent shift. Precision-weighting, therefore, offers a coherent account of the modulatory route while remaining silent on the structural route, which is exhausted by the local repetition-learning rule. The account makes a falsifiable prediction. Directly manipulating response caution, for example, through speed-accuracy instructions or accuracy-emphasizing payoffs, should alter the distractor-absent verification profile while leaving matched-repetition capture and the repetition-learning slope largely stable.

The signature of volatility is recognizable across cognitive domains. In reinforcement learning, volatile contexts raise learning rates and speed the updating of value estimates (Behrens et al., 2007; Piray & Daw, 2020). In anxiety and mood, they reshape how uncertainty itself is tracked and weighted (Pulcu & Browning, 2025). In perception and action, they prompt broader Bayesian-precision adjustments at multiple levels of the inferential hierarchy (Friston, 2009, 2010; C. Mathys et al., 2011; C. D. Mathys et al., 2014; Yu & Dayan, 2005). In perceptual decision-making, they shift commitment thresholds within sequential-sampling models, slowing the rate at which evidence is converted into a response (Forstmann et al., 2016; Ratcliff & McKoon, 2008). The recurring theme is a policy shift. Volatile environments make read-out more conservative, weigh recent evidence more heavily, and demand more before commitment. Stimulus-level processing itself tends to be spared. The present results add visual selection to the list, with one stage-level refinement. The policy shift is visible at the post-selective verification stage in eye movements and at the decision-boundary axis in the manual response. The pre-selective gating stage and the local repetition-learning rule are preserved. Volatility, in this view, is a domain-general modulator of decision policy that reaches selective attention through its read-out stage.

The latent-decision models reinforce this interpretation in a limited, descriptive sense by separating decision-state axes that raw RT cannot separate. The HSSM boundary-only projection estimated a higher decision boundary in high-volatility blocks, giving the modulatory route from the eye-movement data a formal decision-process counterpart. The drift-only projection estimated lower present-trial drift under high volatility (Δ*v* = −0.29), and the full model outperformed boundary-only and drift-only alternatives in PSIS-LOO model comparison. Manual responses, therefore, contain both boundary- and drift-related signals. The repetition-aware ocular result clarifies how the drift component should be interpreted. It is not evidence for a per-trial loss of oculomotor gating, because once the same-location repetition is matched, no per-trial capture difference remains. A complementary VKF dynamic-state comparison reached a compatible architecture: the best aggregate AIC favored a model in which the latent volatility/prediction-error state modulates both drift and boundary. BIC, by contrast, favored a simpler gain-only baseline, and the VKF likelihood is only an approximation to the HSSM Wiener model. We therefore do not describe volatility as boundary-only at the parameter level. Manual responses contain both axes, and the eye-tracking analyses constrain how the drift axis must be read.

Same-location repetition learning showed no detectable difference in slope across volatility regimes on every measure we examined. This pattern converges with classical priming-of-pop-out (Allenmark, Gokce, et al., 2021; Maljkovic & Nakayama, 1994, 1996; Xia et al., 2026), with trial-by-trial dynamic statistical-learning evidence that the priority map is reshaped across consecutive distractor-present trials (Allenmark et al., 2022, 2024; Ferrante et al., 2018), and with accounts in which repeated salient stimuli progressively lose their grip on attention or become easier to suppress (Turatto and Valsecchi 2022; Won et al. 2019; Müller et al. 2009). Statistical-learning accounts predict that sequential regularities update suppression weights regardless of higher-order block structure. The absent slope difference we observe is exactly the pattern those accounts predict. What the present data add is that the repetition-learning rule remains stable under environmental volatility that observers cannot readily report. The structural route through which volatility reaches early selection in this design, therefore, relies on the same learning machinery as in stable environments. Volatility changes the *repetition distribution* that feeds the learning rule, leaving the rule itself untouched.

The present results complement recent volatility work in distractor-bias designs. Qiu et al. (2024) varied volatility in a paradigm with a strong long-term spatial bias in distractor location and showed that volatility modulated the deployment of the learned spatial prior during target selection. Our design deliberately removes the long-term spatial prior, so suppression had to be built solely from short-term repetition. The structural route, therefore, surfaces in our data as a run-position mixture effect, whereas in Qiu et al. (2024) it surfaced as modulation of a learned spatial prior. Taken together, the two studies suggest that volatility reaches the priority map via whatever learned regularity is locally available, while reserving its modulatory route for the readout stage. The present data sit alongside Bogaerts et al.’s (2022) and van Moorselaar and Slagter’s (2019) demonstrations that distractor regularities at different timescales modulate suppression. We extend those findings by isolating volatility, with overall distractor frequency held constant, as the manipulated factor. The pattern also dovetails with stage-aware terminology proposals that recommend separating pre-selective, post-capture, and post-selective measurement targets (Liesefeld et al., 2024; Sauter et al., 2021): the modulatory route is detectable precisely at the post-selective stage that those proposals identify as criterion-driven (Allenmark, Shi, et al., 2021).

Continuing this dissociation will require several extensions to the current design. The retained sample of twenty-one observers gives medium-effect sensitivity; a larger sample would sharpen the matched-repetition capture estimate and the precision with which formal equivalence can be tested at small-effect bounds. The HSSM and VKF analyses identify useful latent-decision axes without uniquely partitioning the full *v*–*a*–*t* space, which is why we treat their convergence with the eye-tracking analyses as the load-bearing claim and the parameter magnitudes as descriptive. Volatility was defined here by the binary presence or absence of the distractor; defining it instead by distractor location or feature would be a more demanding test, especially because filtering distractors defined in a different dimension than the target can follow a partially distinct logic (Sauter et al., 2018). Awareness of the manipulation was poor: only one retained participant answered the post-experiment probe correctly. The pattern, therefore, reflects an implicit adaptation to sequence structure, and an explicit-instruction version of the paradigm would clarify whether observers can deliberately deploy or override the post-selective caution we observed.

A generalisable design principle follows from the structural-route reading. Any volatility manipulation that operates through the transition probability of a binary latent state imposes distinct distributions of consecutive same-state run positions on its two regimes. Whenever the dependent measure is sensitive to run position, and almost every short-term distractor-suppression measure is (Allenmark, Gokce, et al., 2021; Feldmann-Wüstefeld & Schubö, 2016; Maljkovic & Nakayama, 1994, 1996; Xia et al., 2026), the pooled volatility contrast will mix the per-trial volatility effect with the rep-mixture the design itself produces. Pooled contrasts remain valid summaries of behavior under each regime and the right summary for characterizing the lived phenotype of volatility. Mechanistic inference about a per-trial volatility effect, however, requires conditioning on, or partialing out, run position. The same logic applies wherever state-persistence manipulations produce state-dependent trial dynamics, including reward-environment paradigms that vary contingency volatility and cued-task-switching designs that vary switch probability.

Taken together, the present study establishes a stage-specific account of how volatility shapes visual selection. Oculomotor capture is invariant to environmental volatility once the same-location repetition is matched, undercutting the intuition that volatile contexts weaken pre-selective distractor suppression. The genuine volatility signature appears at the post-selective verification stage on distractor-absent trials and is mirrored by a higher latent decision boundary in the manual response, identifying the decision criterion rather than early gating as the locus at which volatility reaches visual selection. The local same-location repetition-learning rule is preserved across regimes; volatility reaches the priority map by reshaping which trial histories the rule encounters, leaving the rule itself untouched. The implications follow directly. Volatility joins the family of domain-general criterion modulators alongside perceptual choice, reinforcement learning, and uncertainty-driven affective regulation and should not be cast as a manipulation that selectively impairs early visual filtering. State-persistence designs, by their construction, recruit a structural route through whatever local learning rule is in play, so mechanistic claims about the volatility effect on attention require designs that condition on or partial out the trial-history mixture; theories of volatility-driven control belong at the readout stage where the present data place them.

## Data availability statement

Data and code are available at https://github.com/msenselab/volatility-shifts-capture-to-caution

## Acknowledgement

This work was supported by a German Science Foundation (DFG) research grant SH 166/10-1 to Z.S and the Sichuan Province Postdoctoral Research Special Funding Project (No. TB2025066) to N.Q. The authors declare no conflict of interest. During the preparation of this work, the authors used ChatGPT and Grammarly AI to polish the language and tidy and optimize Python analysis codes. After using this tool/service, the authors reviewed and edited the content as needed and take full responsibility for the published article.

